# Elevations in plasma proinsulin predict the development of diabetes in NOD mice

**DOI:** 10.1101/2025.07.07.663576

**Authors:** Vriti Bhagat, Jorden Gill, Mélanie Lopes, C. Bruce Verchere

**Author notes:** Corresponding author: C. Bruce Verchere, PhD, BC Children’s Hospital Research Institute, 950 W 28^th^ Avenue, Vancouver, BC, Canada V5Z4H4.

## Abstract

The global incidence of type 1 diabetes (T1D) continues to rise, yet reliable biomarkers for predicting disease onset remain limited. Studies have demonstrated persistent proinsulin secretion in individuals living with T1D, suggesting a processing impairment. Proinsulin is processed into mature active insulin by the prohormone convertases PC1/3, PC2, and carboxypeptidase E. We hypothesized that elevated circulating proinsulin-to-C-peptide (PI:C) ratios precede the onset of diabetes and are associated with reduced expression of PC1/3 in pancreatic beta cells. Non-obese diabetic (NOD) mice were monitored for changes in plasma proinsulin, C-peptide, and beta cell Pc1/3 levels prior to diabetes onset. Female NOD mice that progressed to diabetes exhibited increased plasma proinsulin and PI:C ratios several weeks before the onset of diabetes compared to mice that remained normoglycemic. Plasma proinsulin levels were predictive of diabetes onset, with earlier elevations observed in mice that progressed to disease more rapidly. These increases in plasma proinsulin and PI:C ratios correlated with reduced beta cell Pc1/3 expression. These findings support the potential of plasma proinsulin and PI:C ratios as predictive biomarkers for T1D development and implicate diminished Pc1/3 expression as a possible mechanism underlying impaired proinsulin processing.

## Introduction

Type 1 diabetes (T1D) incidence is increasing worldwide, with a global increase of 3-4% of new cases diagnosed each year^1^. T1D is an autoimmune disorder marked by chronic hyperglycemia resulting from the destruction of insulin-producing pancreatic beta cells^2^. Genetic predisposition plays a significant role in disease risk, particularly through HLA class II haplotypes and polymorphisms in genes related to insulin production and immune regulation^2,3^. While individuals inherit genetic risk for T1D, initiation of autoimmunity in T1D is thought to require an environmental or physiological trigger ^4^. The presence of multiple autoantibodies targeting beta cell proteins is associated with an increased risk of disease progression^5^. Individuals with autoantibodies against two or more beta cell antigens face a 27-70% risk of developing T1D within 10 years^5^; however, not all individuals with autoantibodies will ultimately develop clinical disease^6,7^. This underscores a critical need to identify additional biomarkers capable of accurately predicting which individuals are at the highest risk of progressing to clinical T1D. Such biomarkers would not only improve predictive accuracy but also open opportunities for early therapeutic intervention, potentially delaying or even preventing the onset of T1D.

Insulin is initially synthesized as the precursor protein proinsulin, which is cleaved by the prohormone convertases PC1/3 and PC2, and further processed by carboxypeptidase E (CPE) within the beta cell secretory pathway^8^. Elevated proinsulin secretion has been identified in individuals with T1D^9^ and in NOD mice^10^, serving as a potential marker of beta cell dysfunction and stress. Previous studies suggest that the proinsulin-to-C-peptide (PI:C) ratio may have value as a predictive biomarker for T1D risk^11,12^; however, these investigations have typically involved cross-sectional analyses in autoantibody-positive individuals, without longitudinal monitoring of proinsulin and C-peptide levels prior to disease onset. As a result, the dynamic changes in these biomarkers, and their potential to predict which individuals progress to T1D, remain inadequately understood.

To assess the potential of proinsulin levels as a predictor of T1D, we longitudinally monitored plasma proinsulin and C-peptide levels in NOD mice every two weeks starting at 8 weeks of age and compared the trajectories of mice that progressed to diabetes and those that remained non-diabetic. We also performed immunohistochemistry on pancreatic tissue from NOD mice at various ages. We found that early elevations in plasma proinsulin were predictive of eventual diabetes development in female NOD mice, and that increased plasma proinsulin and PI:C ratio were associated with a decrease in beta cell Pc1/3 levels. These findings support the potential utility of plasma proinsulin and PI:C ratio as non-invasive biomarkers for predicting T1D risk and provide mechanistic insights into beta cell dysfunction preceding clinical onset.

## Methods

### Mouse Studies

Female and male NOD/ShiLtJ (NOD) mice were bred in-house, and age- and sex-matched NOD.Cg-*Prkdc*^*scid*^/J (NOD.Scid) mice were purchased from The Jackson Laboratory (Bar Harbor, ME, USA). Blood samples were collected biweekly via the saphenous vein into EDTA-coated tubes. For terminal studies, blood was obtained through cardiac puncture. Blood samples were centrifuged at 2,000g for 6 minutes at 4°C to isolate plasma. Plasma proinsulin levels were quantified using a Rat/Mouse Proinsulin ELISA kit (10-1232-01, Mercodia), and plasma C-peptide concentrations were determined using a Mouse C-peptide ELISA kit (80-CPTMS-E01, Alpco Diagnostics). Diabetes onset was defined by at least one blood glucose measurement above 20 mM. All animal protocols complied with the Canadian Council on Animal Care guidelines and received approval from the University of British Columbia Animal Care and Use Committee.

### Immunostaining and Image Analysis

Mice were euthanized by isoflurane inhalation under a surgical plane of anesthesia and transcardial perfusion immediately performed with phosphate-buffered saline (PBS) and 4% paraformaldehyde (PFA). The pancreas was then harvested, bisected into head and tail regions, and fixed overnight in 4% PFA at 4 °C. Tissues were subsequently transferred to 70% ethanol and stored at 4 °C. Fixed pancreas tails were embedded in paraffin, and sliced into 5 μm thick sections. Slides were deparaffinized through xylene washes and rehydrated with a series of ethanol washes. Antigen retrieval was conducted in preheated 10 mM citrate buffer (pH 6). Sections were blocked for 1 hour at room temperature using Dako blocking solution (X0909, Agilent). Following washes, tissue sections were incubated overnight at 4 °C with a primary antibody cocktail containing guinea pig anti-insulin (1:4; IR00261-2, Dako), mouse anti-proinsulin (1:50; GS-9A8, DSHB), and rabbit anti-Pc1/3 (1:500; ab220363, Abcam). After washing, slides were incubated for 1 hour at room temperature with a secondary antibody cocktail comprising DAPI (5 mg/mL, 1:1000; D3571, Invitrogen), goat anti-guinea pig, anti-mouse, and anti-rabbit antibodies (each 1:250; Invitrogen). Sections were then washed and mounted in ProLong™ Gold Antifade Mountant (P36930, Invitrogen). Tiled fluorescent images were acquired using a 10x objective on an Olympus BX61 microscope, with consistent exposure settings applied across all sections. For each mouse, 1-2 non-consecutive pancreatic sections were imaged. Images were analyzed using QuPath version 0.5.0 as previously described^13,14^. In brief, islets were segmented, nuclei were detected, and insulin-, proinsulin-, and Pc1/3-positive cells were identified based on cell mean intensity thresholds. Mean fluorescence intensities were quantified on a per-cell basis, with background fluorescence subtracted using adjacent non-islet tissue regions.

### Statistical Analysis

Statistical analyses were performed using GraphPad Prism 10. Datasets with normal distribution were analyzed by Student’s t-test or one-way ANOVA. Non-normally distributed datasets were analyzed by Mann-Whitney U test or Kruskal-Wallis test. Statistical significance is indicated in the figures as \**p*<0.05, \*\**p*<0.01. All data are presented as mean ± SEM.

### Data and Resource Availability

Data generated in the current study are available upon reasonable request from the corresponding author.

## Results

### Plasma proinsulin and proinsulin-to-C-peptide is elevated several weeks prior to the onset of diabetes

To explore whether plasma proinsulin levels differ between non-diabetic and diabetic mice prior to the development of autoimmune diabetes, female and male NOD mice were grouped based on their diabetes status at the end of the 30-week study period. Age- and sex-matched NOD.Scid mice served as controls. Body weight was comparable between female and male NOD mice and NOD.Scid controls throughout the study (Fig. 1A and Supplementary Fig. 1A). Blood glucose >20 mM was considered diabetic, with onset beginning at 14 weeks of age for female NOD mice and 20 weeks of age for male NOD mice. The median age of diabetes onset for female NOD mice was 18.5 weeks, while for male NOD mice it was 22 weeks (Fig. 1B and Supplementary Fig. 1B). Consistent with previously reported figures in the literature^15^, the incidence of diabetes was 86.6% for female NOD mice and 20% for male NOD mice in this cohort (Fig. 1C and Supplementary Fig. 1C). Blood samples were collected every two weeks to monitor plasma proinsulin and C-peptide levels via ELISA. Notably, female NOD mice that developed diabetes showed 1.73-fold higher plasma proinsulin levels and 1.42-fold higher PI:C ratios 7-10 weeks prior to diabetes onset, compared to female NOD mice that remained non-diabetic throughout the study. These differences in PI:C ratios persisted until the end of the study, while proinsulin levels remained elevated up to one week before diabetes onset, after which they declined likely due to beta cell loss (Fig. 1D-F). In contrast, no such trends were observed in male NOD mice (Supplementary Fig. 1D-F).

**Figure 1.**
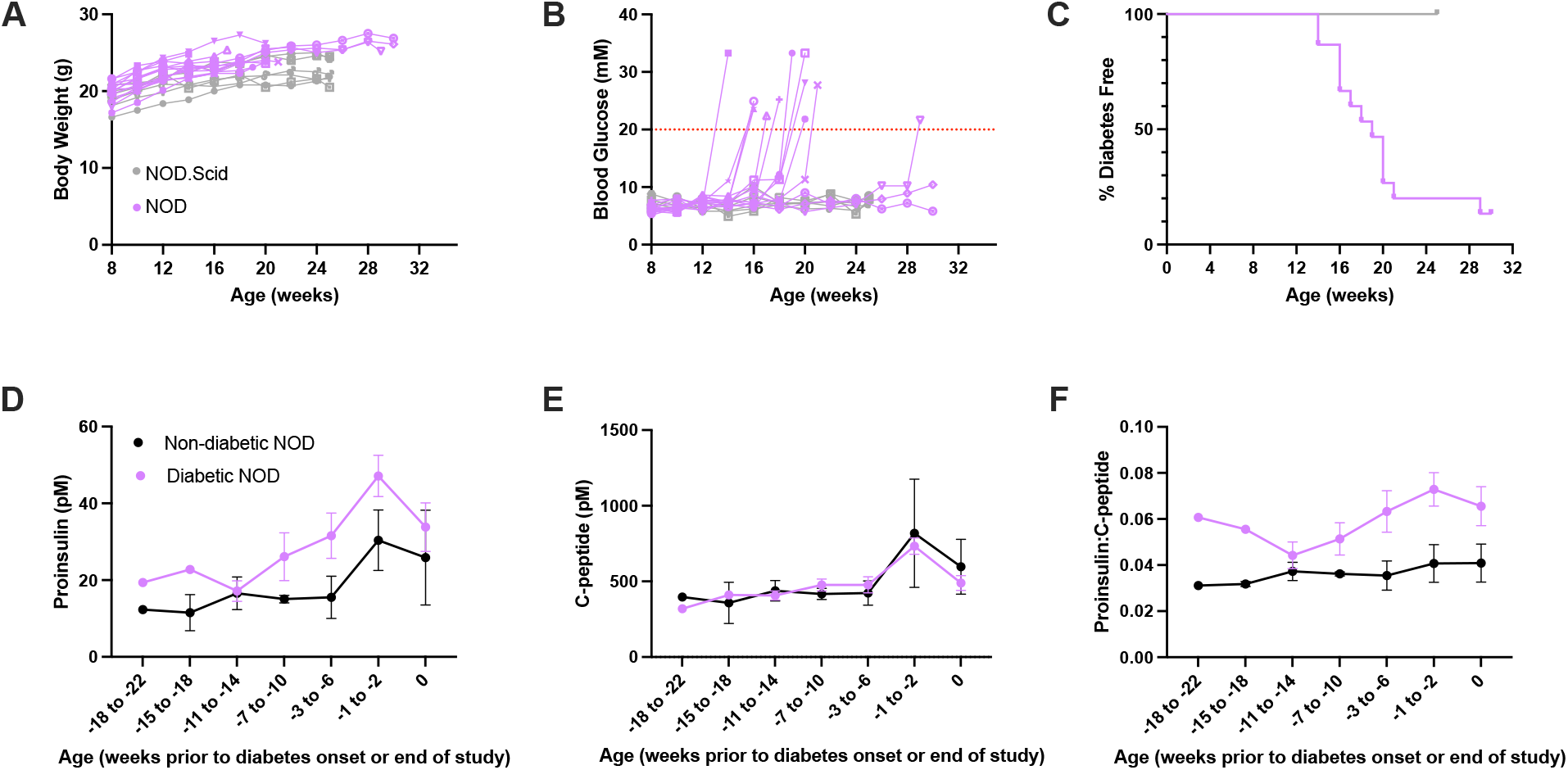
Plasma proinsulin and proinsulin-to-C-peptide ratio is elevated several weeks prior to diabetes onset. (A) Body weight of female NOD (n=14) and NOD.Scid (n=6) mice. (B) Random blood glucose measurements. Diabetes onset was defined as at least one blood glucose measurement above 20 mM as indicated by the red dotted line. (C) Diabetes incidence in NOD compared to NOD.Scid mice. (D) Random plasma proinsulin, (E) C-peptide, and (F) proinsulin:C-peptide levels in diabetic vs non-diabetic mice. Error bars represent mean ± SEM.

### Proinsulin levels predict diabetes development in female NOD mice

To assess the potential of elevated proinsulin as a predictive biomarker for diabetes development, female NOD mice were categorized into three groups: early-onset diabetic, late-onset diabetic, or non-diabetic. Early-onset was defined as diabetes occurring before the median onset age of 18.5 weeks (Fig. 2A), while late-onset was defined as diabetes development after 18.5 weeks (Fig. 2D). Male NOD mice were separated into 2 groups, diabetic and non-diabetic, due to the low number of male diabetic mice. Notably, female NOD mice that exhibited elevated plasma proinsulin levels early, at 8 weeks of age, developed diabetes prior to the median onset age (Fig. 2B). At 8 weeks, plasma proinsulin levels were 2.52-fold higher in the early-onset diabetic females compared to the late-onset and non-diabetic group. Further, although the late-onset diabetic group displayed no difference in plasma proinsulin levels at 8 weeks of age, proinsulin levels were 2.45-fold higher in this group at 16 weeks compared to mice that remained non-diabetic throughout the study (Fig. 2E). Additionally, PI:C ratios trended upwards in both groups at these time points (Fig. 2C and F). These trends were not observed in male NOD mice (Supplementary Fig. 2A-B). Thus, in female NOD mice, elevated plasma proinsulin levels predicted which mice would develop diabetes, occurring several weeks before the actual onset of diabetes in both early- and late-onset groups.

**Figure 2.**
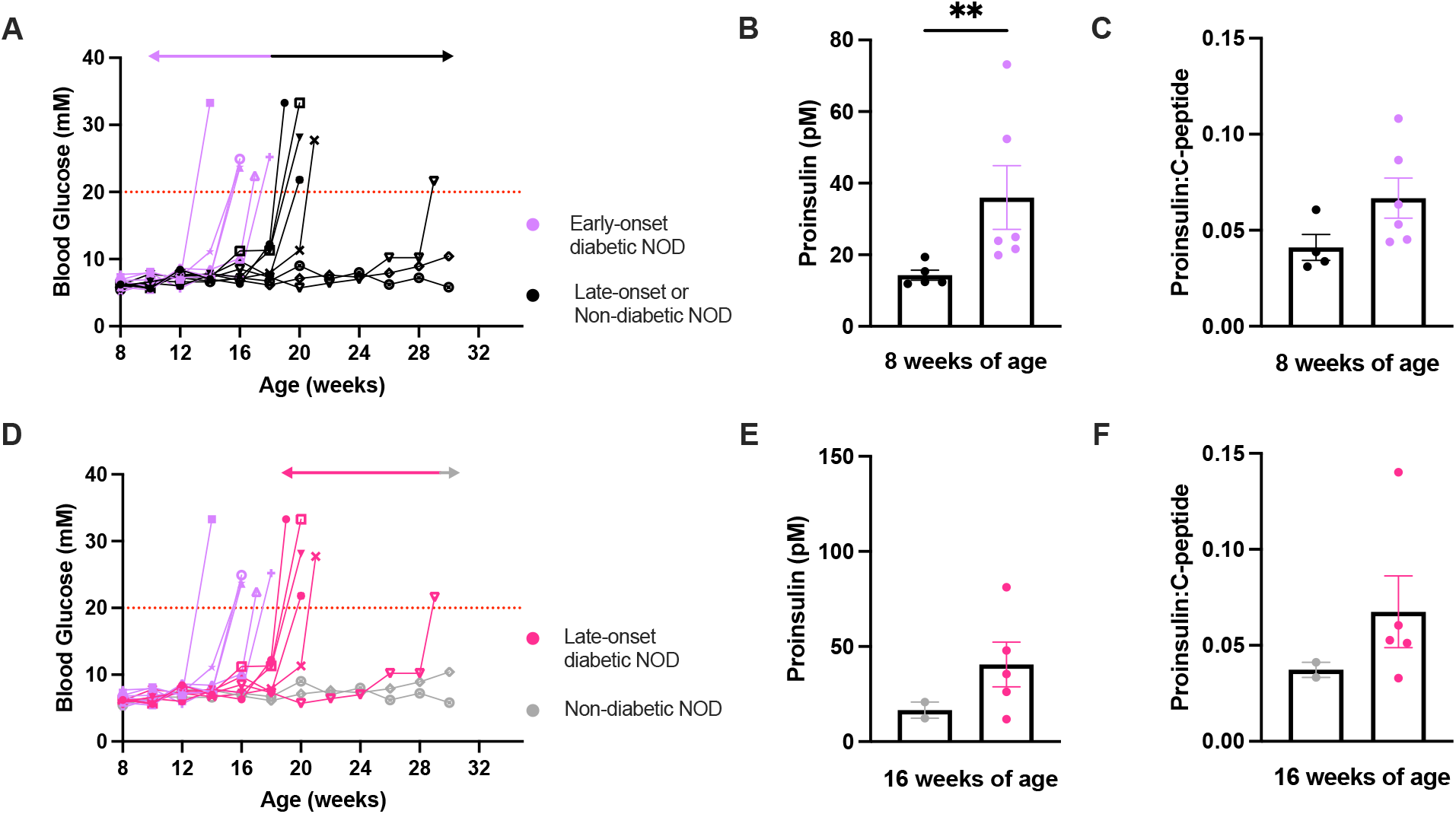
Elevations in plasma proinsulin predict diabetes development. (A,D) Female NOD mice were divided into three groups: early-onset diabetic NOD mice developed diabetes prior to the median onset age of 18.5 weeks, late-onset diabetic NOD mice developed diabetes after 18.5 weeks, and non-diabetic mice remained normoglycemic throughout the study. (B) Plasma proinsulin levels and (C) proinsulin:C-peptide at 8 weeks of age in mice that developed diabetes before the median onset age compared to mice that did not develop diabetes. (E) Plasma proinsulin levels and (F) proinsulin:C-peptide at 16 weeks of age in mice that developed diabetes after the median onset age compared to mice that did not develop diabetes. Error bars represent mean ± SEM. \*\**p* < 0.01.

### Beta cell Pc1/3 levels decrease as female NOD mice progress towards diabetes

To investigate potential mechanisms underlying the elevation of plasma proinsulin in pre-diabetic NOD mice, the tail of the pancreas was harvested from a separate cohort of female NOD mice at 6, 8, 12, and 16 weeks of age, all prior to diabetes onset (Supplementary Fig. 3B). These pancreas sections were analysed by quantitative immunohistochemistry for insulin, proinsulin and Pc1/3 levels. Of note, female NOD mice showed a 1.84-fold reduction in beta cell Pc1/3 levels between 6 and 16 weeks of age (Fig. 3A-B). Of note, this reduction did not correlate with a decrease in the number of beta cells expressing Pc1/3 (Fig. 3C), a change in the colocalization of insulin and Pc1/3 (Fig. 3D), or an increase in the number of non-beta cells expressing Pc1/3 (Fig. 3E). The reduction in Pc1/3 levels within beta cells was associated with an 11.32-fold increase in plasma proinsulin levels (Supplementary Fig. 3C).

**Figure 3.**
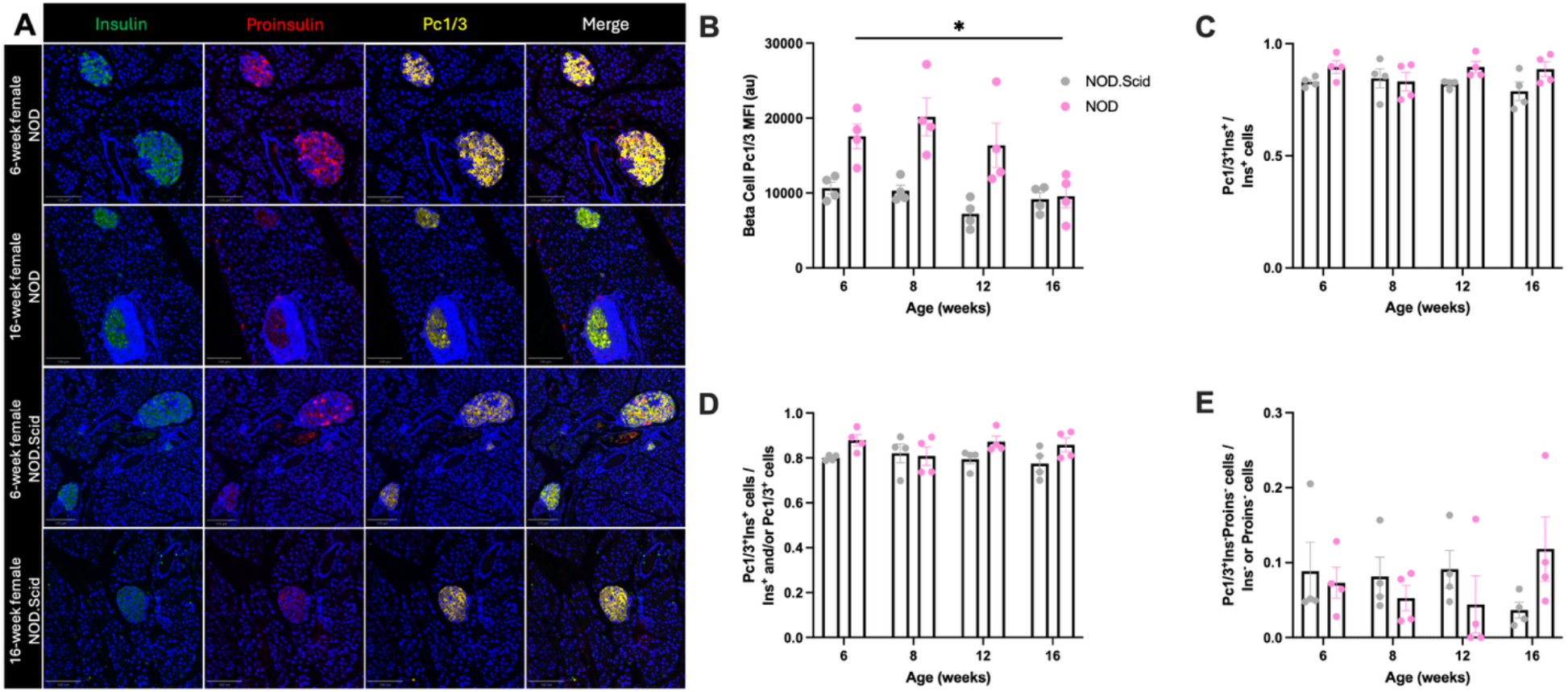
Beta cell Pc1/3 levels decrease as female NOD mice progress towards diabetes. (A) Representative images from 6- and 16-week-old female NOD and NOD.Scid mice pancreas tail sections stained for insulin, proinsulin, and Pc1/3 expression. Scale bar = 100 µm. (B) Pc1/3 mean fluorescent intensity in Ins^+^Pc1/3^+^ cells. (C) Proportion of Ins^+^ cells that express Pc1/3. (D) Colocalization of insulin and Pc1/3 expression. (E) Proportion of Pc1/3^+^Ins^-^ cells. n = 4 per age and strain. 1-2 non-consecutive sections were analyzed per mouse. Error bars represent mean ± SEM. \**p* < 0.05. MFI, mean fluorescent intensity; au, arbitrary units.

## Discussion

Over the years, studies have demonstrated that proinsulin levels rise prior to the onset of T1D^11,12,16,17^ and remain elevated at onset^9,14,18–20^ in both humans and mouse models^10^. While these earlier studies suggest that elevated proinsulin and/or PI:C ratios could have value as predictive biomarkers of T1D, they have not longitudinally tracked at-risk individuals to assess whether proinsulin levels could serve as a predictive biomarker for disease development. In this study, we show that elevated proinsulin levels can predict ensuing diabetes onset in the well-established NOD mouse model of T1D.

A previous study reported that proinsulin levels and the proinsulin-to-insulin ratio increased with age in pre-diabetic mice^10^. Here we extend these findings by stratifying mice based on diabetic outcome and longitudinally tracking plasma proinsulin levels and PI:C ratio. We found that mice that ultimately developed diabetes exhibited significantly elevated levels of both markers up to 10 weeks prior to disease onset compared to non-diabetic controls, supporting the idea that beta cell prohormones may be predictive biomarkers in T1D^21^. The PI:C ratio remained elevated at disease onset, whereas absolute proinsulin levels declined at the time of onset, likely reflecting beta cell loss. These results support previous reports that beta cell dysfunction precedes clinical onset of human T1D^22^. Several factors may contribute to increased circulating proinsulin during this prediabetic phase, including endoplasmic reticulum (ER) stress in beta cells^10^, impaired proinsulin conversion^19^, beta cell dedifferentiation^21^, and/or increased proinsulin secretion in response to elevated insulin demand^16^. Future studies would benefit from longitudinal analyses in human cohorts, particularly among individuals with a family history of T1D, both those who are autoantibody-negative and autoantibody-positive, to better characterize these biomarker fluctuations in human disease progression.

A previous study examining a cohort of autoantibody-positive individuals reported that those who progressed to T1D within 12 months had significantly higher plasma proinsulin levels compared to those who remained non-diabetic^11^. Our study demonstrates that early elevation of plasma proinsulin is closely associated with the timing of autoimmune diabetes onset. Mice that developed diabetes at a younger age, before 18.5 weeks, consistently showed elevated proinsulin levels early on at 8 weeks of age. In contrast, mice that developed diabetes later, after 18.5 weeks, did not exhibit elevated proinsulin at 8 weeks, but levels rose markedly by 16 weeks, shortly before disease onset. These findings underscore the potential of elevated proinsulin as an early and informative biomarker for predicting the development of T1D. In our cohort, while all mice with high proinsulin levels eventually developed diabetes, a few mice with lower levels in the late-onset group also progressed to disease, suggesting that proinsulin may be more effective as part of a multi-biomarker approach rather than as a standalone predictor. Indeed, prior research has proposed that combining autoantibody status with proinsulin measurements could improve risk prediction^12^, given that not all autoantibody-positive individuals progress to diabetes^6,7^. Other potential biomarkers that could be used in combination with autoantibody status and proinsulin levels for T1D prediction include genetic risk score^23^, T cell biomarkers^24^, and other beta cell biomarkers^21^ such as pro-islet amyloid polypeptide (proIAPP)^25^. Of note, PI:C ratio was not predictive in our study, though this may be attributable to the limited sample size, since an upward trend in the PI:C ratio was observed prior to disease onset.

Despite numerous studies demonstrating elevated proinsulin levels in T1D^9–12,14,16–20^, the underlying mechanisms remain unclear. Some research has shown reduced islet PC1/3 levels in pancreatic tissues from individuals living with T1D^19,26^. More recently, a study examined PC1/3 expression in both beta and alpha cells from donor T1D pancreatic tissue^14^. In our study, we specifically investigated Pc1/3 levels within insulin-producing beta cells in pre-diabetic NOD mice. We found that as NOD mice progress toward diabetes, Pc1/3 expression in beta cells was significantly reduced, whereas the proportion of Pc1/3+Ins+ cells, the degree of colocalization between Pc1/3 and insulin, and the number of Pc1/3+ non-beta cells remained unchanged. This finding suggests that the observed reduction in Pc1/3 occurs within individual beta cells rather than due to a loss of Pc1/3-expressing beta cells. The observed decline in beta cell Pc1/3 expression likely contributes to the elevated circulating proinsulin levels observed during disease progression. The mechanism underlying reduced Pc1/3 expression in beta cells prior to T1D onset remains unknown but could reflect beta cell stress and dysfunction in the early stages of autoimmunity.

One important consideration in this study is that differences in proinsulin and PI:C were observed exclusively in female NOD mice, with no corresponding changes detected in males. As male NOD mice have a lower incidence of diabetes^15^, this finding is in keeping with impaired proinsulin processing being a consequence and/or contributor to beta-cell directed autoimmunity and the pathogenesis of T1D^21^. The mechanisms underlying the relative resistance of male NOD mice to diabetes, despite the presence of insulitis at early ages^27^, remain poorly understood.

In summary, our findings demonstrate that both plasma proinsulin levels and PI:C ratio are elevated several weeks prior to the onset of diabetes and that elevated proinsulin levels can predict T1D in female NOD mice. Our study provides new insights into the mechanisms that may underlie increased proinsulin levels in T1D and supports the potential utility of proinsulin as a biomarker for predicting disease onset. Further investigation into the factors contributing to elevated plasma proinsulin is warranted to better understand beta cell dysfunction during the progression of T1D.

## Supporting information

Bhagat 2025 Supplementary Material

## Acknowledgements

The authors would like to thank Drs. Galina Soukhatcheva and Lei Dei (University of British Columbia) for technical assistance, the BC Children’s Hospital Research Institute core facilities for histology and microscopy, and the University of British Columbia modified barrier animal facility.

## Author Contributions

V.B. contributed to study conceptualization, experimental design, investigation, formal analysis, interpretation, and wrote the manuscript. J.G. contributed to study investigation and formal analysis. M.L. contributed to study investigation. C.B.V. contributed to study conceptualization, experimental design, interpretation, reviewed and edited the manuscript, and supervised the project.

