## Supplementary material for "Elevations in plasma proinsulin predict the development of diabetes in NOD mice": Bhagat 2025 Supplementary Material

*Corresponding author:

C. Bruce Verchere, PhD

Vriti Bhagat ORCiD ID: 0000-0001-9423-6759

Mélanie Lopes ORCiD ID: 0009-0001-5665-4826

Bruce Verchere ORCiD ID: 0000-0002-9262-0586

### Supplementary Material

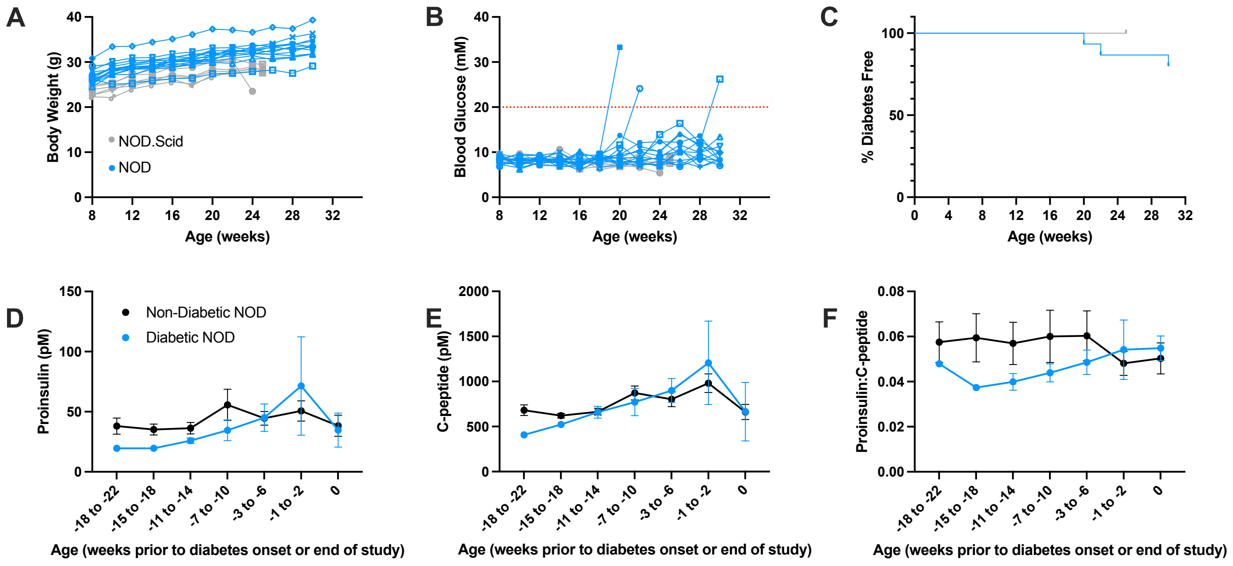


**Supplementary Figure 1. Plasma proinsulin and proinsulin-to-C-peptide ratio is comparable between diabetic and non-diabetic male NOD mice.** (A) Body weight of male NOD (n = 15) and NOD.Scid (n = 6) mice. (B) Random blood glucose measurements. Diabetes onset was defined as at least one blood glucose measurement above 20 mM as indicated by the red dotted line. (C) Diabetes incidence in NOD compared to NOD.Scid mice. (D) Random plasma proinsulin, (E) C-peptide, and (F) proinsulin:C-peptide levels in diabetic vs non-diabetic mice. Error bars represent mean ± SEM.

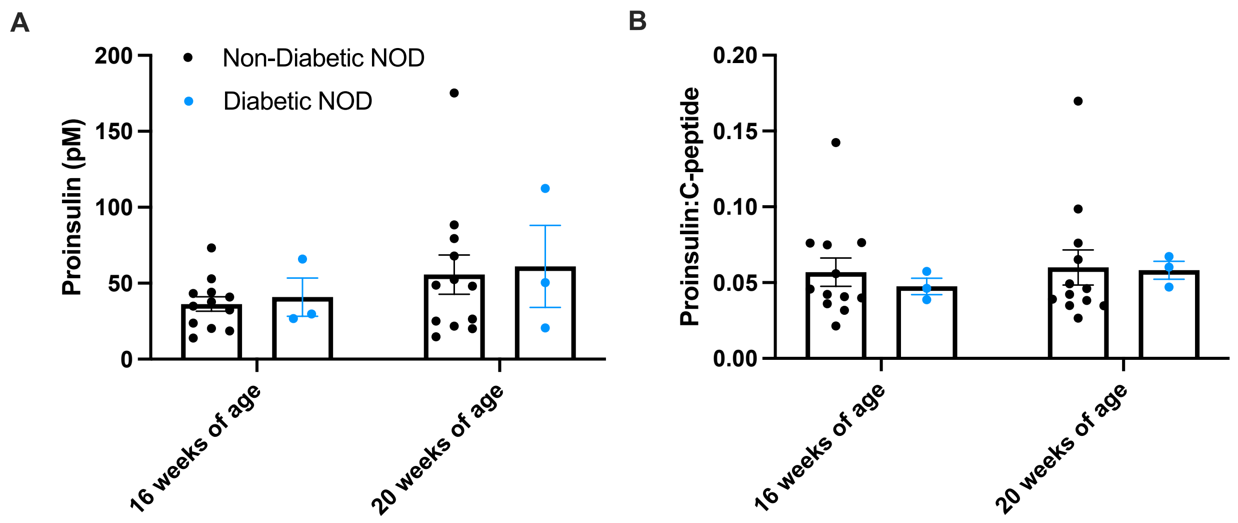


**Supplementary Figure 2. Plasma proinsulin does not predict diabetes development in male NOD mice.** (A) Plasma proinsulin levels and (B) proinsulin:C-peptide in male NOD mice that developed diabetes vs those that did not. Error bars represent mean ± SEM.

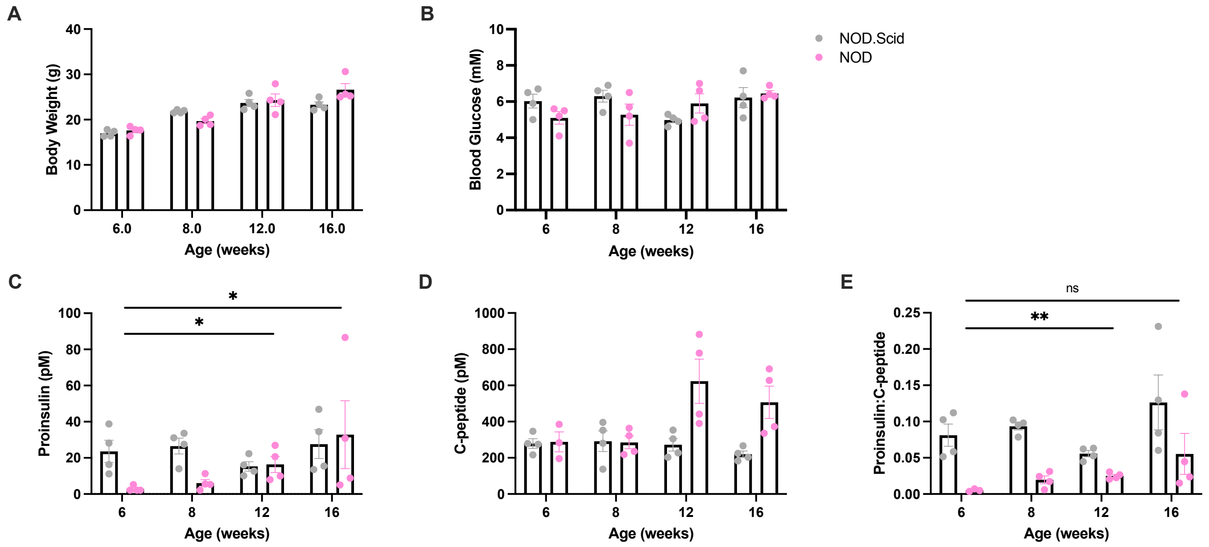


**Supplementary Figure 3. Plasma proinsulin and proinsulin-to-C-peptide ratio increases as female NOD mice progress towards diabetes.** (A) Body weight of female NOD and NOD.Scid mice. 4-hour fasted (B) blood glucose, (C) plasma proinsulin, (D) plasma C-peptide, and (E) proinsulin:C-peptide. n = 4 per age per strain. Error bars represent mean ± SEM. *$p < 0.05$, **$p < 0.01$. ns, not significant.
